# A ready-to-use logistic Verhulst model implemented in R shiny to estimate growth parameters of microorganisms

**DOI:** 10.1101/2022.07.29.501982

**Authors:** Marc Garel, Lloyd Izard, Marthe Vienne, David Nerini, Badr Al Ali, Christian Tamburini, Séverine Martini

## Abstract

In microbiology, the estimation of the growth rate of microorganisms is a critical parameter to describe a new strain or characterize optimal growth conditions. Traditionally, this parameter is estimated by selecting subjectively the exponential phase of the growth, and then determining the slope of this curve section, by linear regression. However, for some experiments, the number of points to describe the growth can be very limited, and consequently such linear model will not fit, or the parameters estimation can much lower and strongly variable. In this paper, we propose a tools to estimate growth parameters using a logistic Verhulst model that take into account the entire growth curve for the estimation of the growth rate. The efficiency of such model is compared to the linear model. Finally, the novelty of our work is to propose a “Shiny-web application”, online, without any programming or modelling skills, to allow estimating growth parameters including growth rate, maximum population, and beginning of the exponential phase, as well as an estimation of their variability. The final results can be displayed in the form of a scatter plot representing the model, its efficiency and the estimated parameters are downloadable.

## Introduction

Understanding the fundamental principles that underpin the rates of growth and reproduction of organisms is of central ecological importance, ultimately affecting long-term evolutionary trajectories of populations and communities. Under variable conditions (i.e. temperature, medium composition, nutrients availability, oxygen) metabolic, biochemical, and physiological processes can affect the growth of an individual, including single cells (Kempes et al., 2012). Furthermore, gene expression in microorganisms is known to be intimately coupled to the growth state of the cell (Scott and Hwa, 2011). For micro-organisms, the growth is non-linear over time and is defined by three successive phases: the lag, the exponential and the stationary phase. During this increase of micro-organisms, the rate at which the number of organisms in a population increases is defined as the growth rate. Usually, in microbiology, the growth rate is a parameter estimated by defining subjectively the exponential phase in the curve and then this part’s linearity (in logarithmic scale) is used to estimate the slope by linear model. According to Zwietering et al. (1990), a better method is to describe the growth of micro-organisms, under different biotic and abiotic conditions (i.e as temperature, pH, salinity and nutrient concentration), using a growth model.

Mathematical models provide tools widely used, for years, to describe growth of microorganisms. In food microbiology, these models allow predicting the shelf life of a food product. This approach allows detecting the critical paths of the production process and optimizing the production and distribution chain (Zwietering et al., 1990). In the environmental field, models can allow finding optimal growth parameter from a new isolated strain (Martini et al., 2013), or describing the behavior of microorganisms under different biotic and abiotic condition (i.e Eichinger, Kooijman, et al. (2009), Eichinger, Poggiale, et al. (2011), and Garel, Panagiotopoulos, et al. (2021)). Numerous models, including Verhulst (1845, 1847), Gompertz (1825), Richards (1959), Schnute (1981), or Stannard et al. (1985) models are applied to adjust observational datasets in order to estimate growth parameter and to predict bacterial growth over time Zwietering et al. (1990). Monod (1949) growth model is another empirical model to describe microbial growth according to substrate concentration. It differs from other models, previously cited, since it is applied for a constant concentration of substrate, introducing the concept of limiting nutrients (Lobry et al., 1992).

The purpose of this paper is to describe a “Shiny”-web application, built with cran R framework, in which a mathematical model, based on the Verhulst model, is embedded to describe the growth of microorganisms. The objective of such approach is not new in term of model applied in microbiology. However, thanks to the development of interactive web-applications, statistics and models are easily available to the microbiology community. This pluridisciplinary (merging modelling and microbiology) approach will lead to new practices easily-applied to determine growth rate of microbial populations with repeatability and transparency in the methodology. We demonstrate that modelling microbial growth is more efficient to estimate growth parameters even with few data points, and less variable depending on the sampling or user.

## Material and methods

### Web application design

The application is accessible at https://hpteam.shinyapps.io/logistic_microbio/, and was entirely designed with GNU R (R Core Team, 2017). The packages used are: ‘stats’ for logistic Verhulst, ‘quantreg’ for quantile regression and 95% confidence interval estimation, and ‘shiny’ for building the web application. Currently the application is hosted on the server https://www.shinyapps.io/, it is available from any computer (independent of the computer’s OS) with an internet access and a web browser. This web application is also available offline, downloading and running into cran R installing required packages with the source available at the following link: https://doi.org/10.34930/DC1DAF1C-09E3-4829-8878-91D0BF0E643E.

The web application includes four mains panels (Figure 2). Firstly, a panel “Upload data” allows uploading data by proposing different types of vector separators (tabulation, comma, Semicolon) and a choice of decimal markers (Figure 1). The file containing data must be in text or csv format. A demo dataset is already available for download.

**Figure 1.**
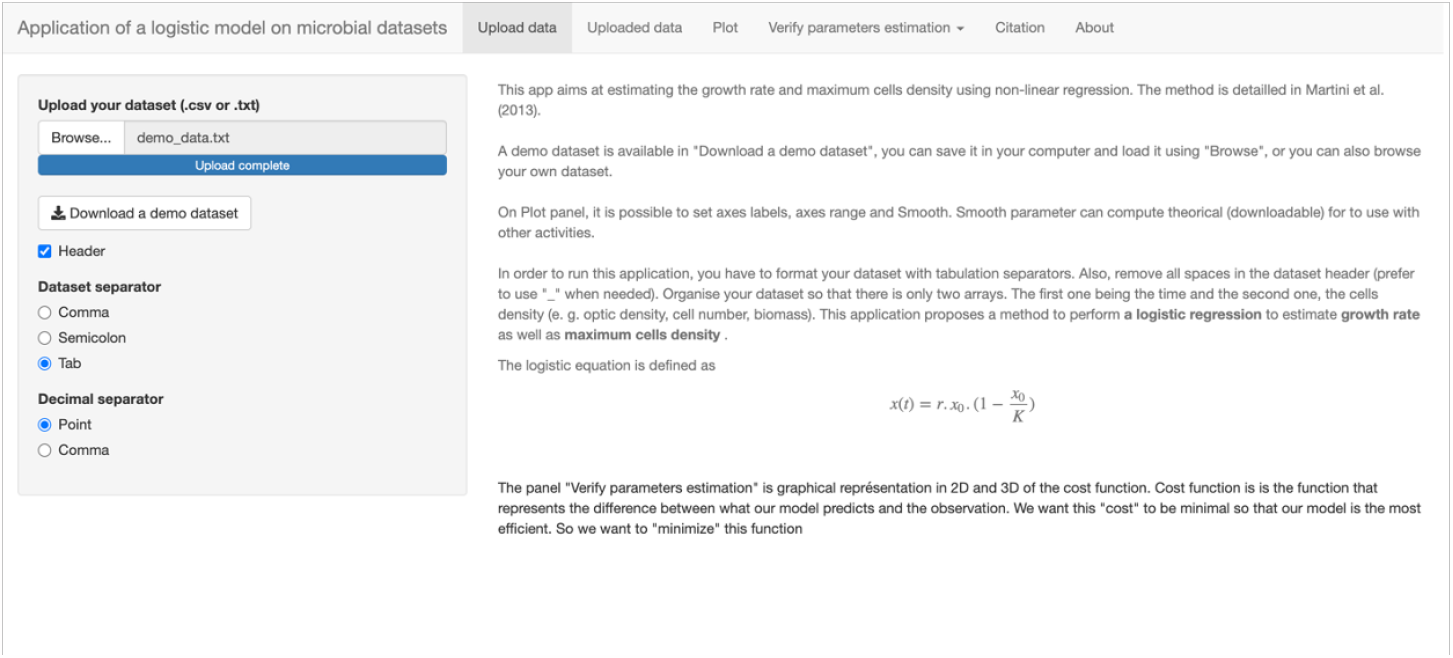
First panel “Upload data” of the web application https://hpteam.shinyapps.io/logistic_microbio/.

**Figure 2.**
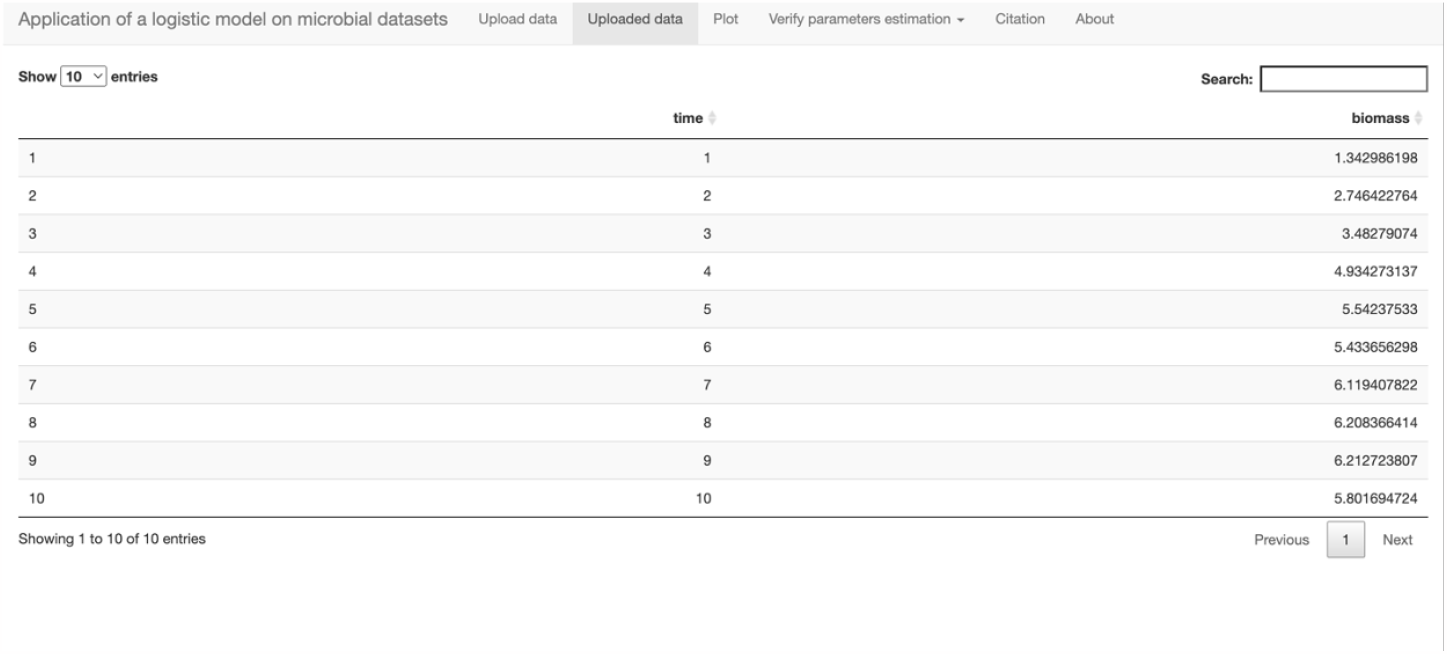
Second panel “Uploaded data” of the application. Dataset is displayed in an interactive dataframe. In this panel dataset can be sorted by entries.

Then, the second panel “Uploaded data” allows viewing the dataset as a dataframe in an interactive table where it is possible to sort data by different variables (Figure 2).

The third panel “Plot” allows viewing the data as a scatter plot Figure (Figure 3). On this tab, it is possible to give a label to each axis, adjust the boundary of each axis and adjust the smoothness of the theoretical curve.

**Figure 3.**
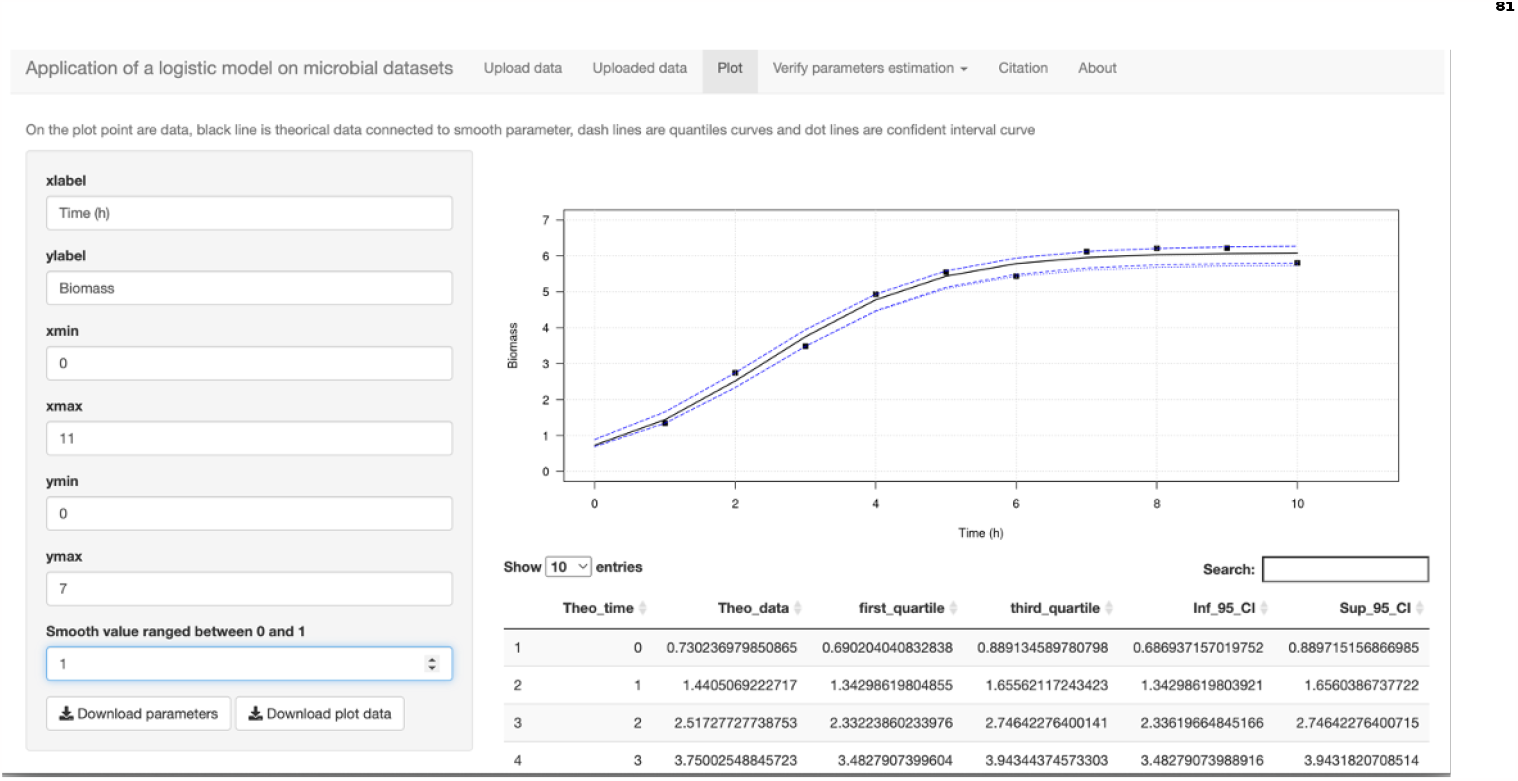
Third panel “Plot” of the application. This panel displays the plot of observed data (dark bullet), fitted data (black line), first and third quartiles (blue dashed line) and 95% confidence interval (blue dot line). The plot can be customized by option : axis label (x and y label), scale of axis (x and y limit) and the smooth of the fitted data. The estimated parameters and fitted curves are downloadable as text file.

By default, the model generates as much theoretical data (data re-estimated by the model) as observed data. This smoothness parameter influences the shape of the curve, the more the “smooth” parameter tends towards 0 the more the curve will have a rounded aspect. However, this parameter has no influence on the estimation of the growth. All results are downloadable in *.txt format. The data set is plotted with model applied on it and the estimated parameters. In this panel, axis label and axis scale are custmomizable and the smooth of the model is adjustable (Figure 3).

Finally, the fourth panel “Verify parameters estimation” allows appreciating the quality of the model using a graphical representation of the Residual Sum of squares (RSS) cost function in 2 or 3 dimensions (Figure 4). Every displayed plots and estimated parameters can be downloaded.

**Figure 4.**
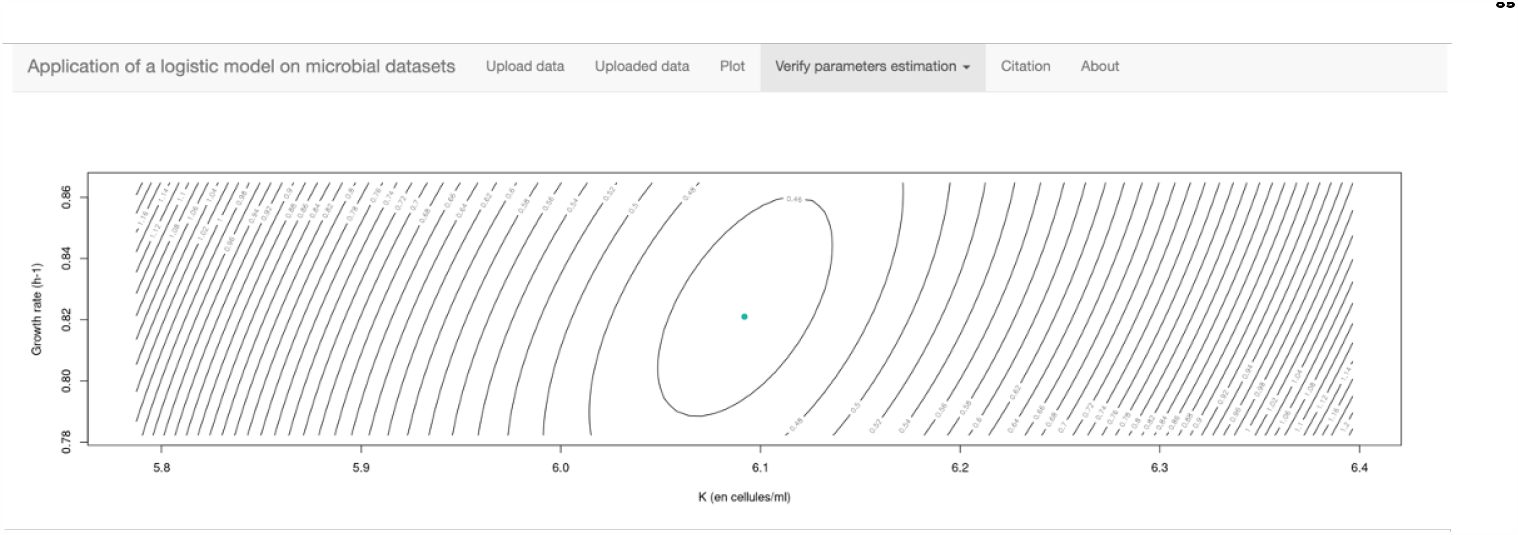
Fourth panel “Verify parameters estimation”. Contour plot showing isoline of the minimisation of growth rate as a function of the maximum of cells (K). The green dot is the minimum of the cost function.

### Model description

The growth of an organism is defined as the variation of the size of the population over time. In our case, growth curve is modelized through a non-linear logistic model described by (Verhulst, 1938). The model is defined by an ordinary differential equation:

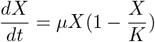

where *X* is the number of individuals, *μ* is the growth rate of the population and *K* the carrying capacity. In microbiology, the growth of prokaryotes is classically characterized by three different phases: a latent phase corresponding to the beginning of the exponential growth. Thereafter, prokaryotes will grow exponentially until they reach a maximum in growth velocity, which will be marked by a decrease of the growth rate to *μ*= 0. This phase is called the stationary phase, which is often due, in the case of batch culture, to a limitation of a substrate, or a nutritive salt involving the population density to remain constant (K). The growth curve thus describes a sigmoid (Zwietering et al., 1990).

The analytical solution of the equation is

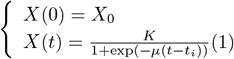

Equation (1) is a sigmoid, a strictly increasing function with time that depends of 3 parameters: *μ*, K, *X*_0_. The experiment provides n pairwise observations (*t*_1_, *X*_1_), …, (*t*_*n*_, *X*_*n*_) which will be used to estimate these parameters. The estimation is achieved when minimizing the following Residual Sum of Squares (RSS).

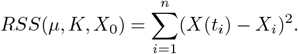

Which computes the squared distance between the observations *X*_*i*_ and model values *X*(*t*_*i*_). As there is no analytical solutions to this non-linear regression problem a numerical iterative Gauss-Newton method is used to estimate the parameters.

For the linear model, traditionally used in microbiology, the growth parameter is estimated by deciding subjectively which part of the curve is approximately linear (in log scale) and then determining the slope of this curve section (Zwietering et al., 1990).

### Model improvement

To test the robustness of the two models (linear and logistic Verhulst) to describe the growth of microorganisms, we created a set of theoretical growth curves using the GNU R statistical software (R Core Team, 2017) representing different cases of study from experimentation. The theoretical curves were constructed with a logistic Verhulst model whose parameters are: experimental time = 10 hours, *μ*_*max*_ =1 *h*^−1^ and *K* as the maximum population density = 6. In addition, a Gaussian noise is added to simulate variability between each measurement point, which is very common in experimentation. Then six experiments are simulated with a decreasing number of points: 100, 75, 50, 25, 10 and 6 points (Figure 5). For each curve the growth parameters is estimated, both with a logistic Verhulst model and with a linear model, classically used in microbiology, by linearizing the data beforehand. For the linear model, the exponential phase was estimated subjectively. The sensitivity of each estimation method was tested with a bootstrap (random re-sampling with discount) of 1000 simulations.

**Figure 5.**
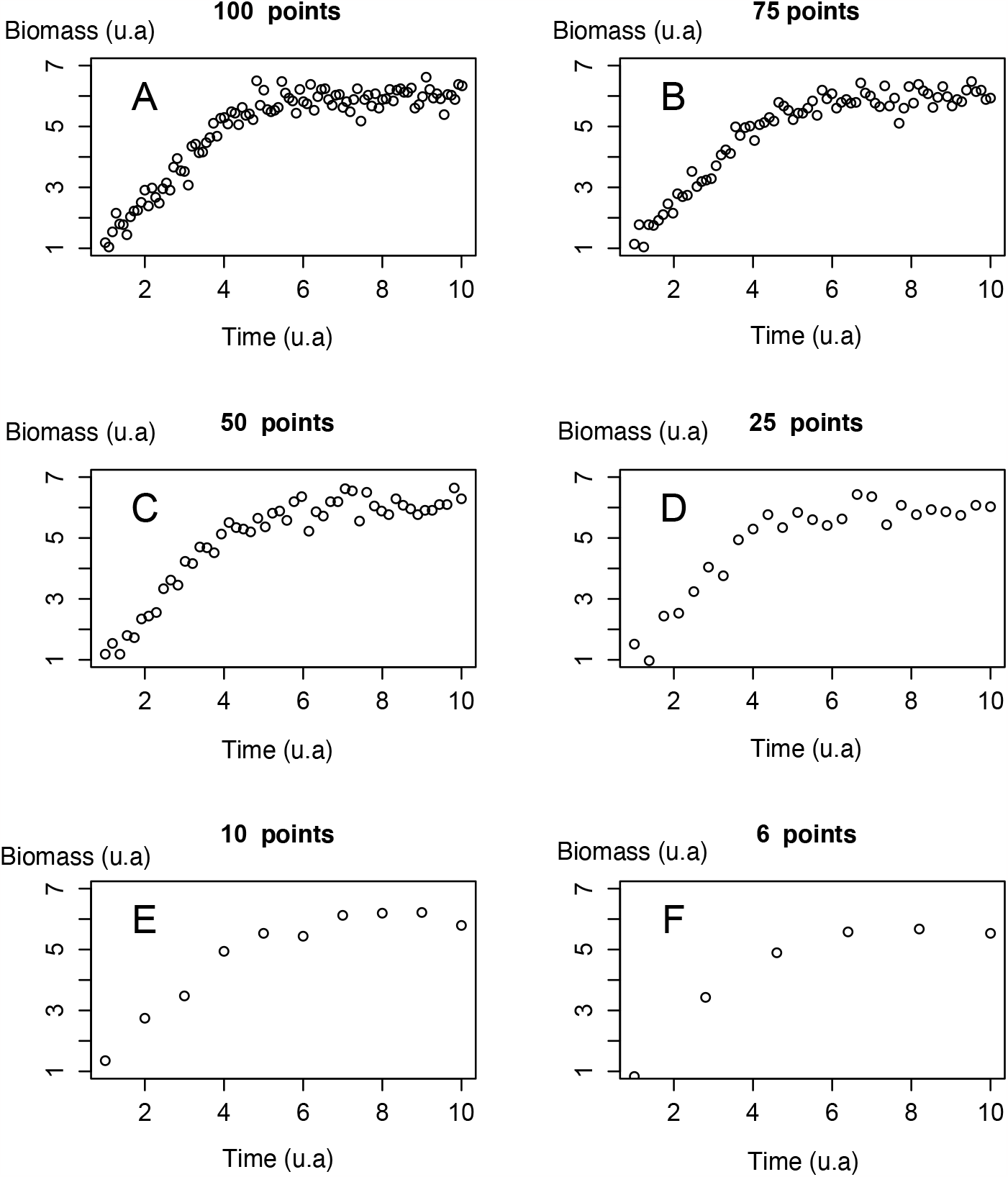
Simulation of logistic growth with different number of measurements to mimic variability in experiments. The data follow a logistic distribution with gaussian noise. The number of points in each curve decreases by random resampling. Biomass is in arbitrary units (u.a.). Time is an arbitrary unit (u.a.) depending to the experimental design.

## Results and Discussion

### Model simulation

The Figure 5 displays six curves simulated by downgrading the number of measured points, from 100 to 6 points, to mimic different cases of experiments. Each succession of points represents the theoretical growth of microorganisms over time. Biomass can be obtained according to different classical methods in microbiology such as the cells number obtained by flux cytometer or microscopy or even by optical density. According to theses curves, growth rate are estimated by bootstrapping.

### Comparison of linear and logistic Verhulst models to estimate growth parameters

To compare the linear and logistic Verhulst models efficiency, we focus on the growth rate parameter *μ*_*max*_, widely used in microbiology in the characterization of new bacterial strains. The following results compare the growth rates’ distribution for various numbers of sampled points ranging from 100 to 6, and using two mathematical approaches: i) estimation of growth parameters using the logistic Verhulst model; ii) estimation of the growth rate by the linear model applied on the exponential phase of growth after linearization (log scale).

In Figure 6, and Table 1 the growth rate parameters estimated using the logistic Verhulst method are close to 1 *h*^−1^ (median values between 0.86 and 1.06 *h*^−1^), corresponding to the growth rate *μ*_*max*_=1 provided as the input in the model. The growth rates *μ*_*max*_ estimated by linear model are about a half of this value (median value between 0.32 and 0.48 *h*^−1^). For both methods, the estimation of the growth rate decreases significantly (wilcox test *p* − *value <* 0.05) for curves built with only 6 points from a median equal to 1.02 to 0.86 *h*^−1^ and from 0.43 to 0.32 *h*^−1^, respectively for logistic Verhulst model and linear model model.

**Table 1.**
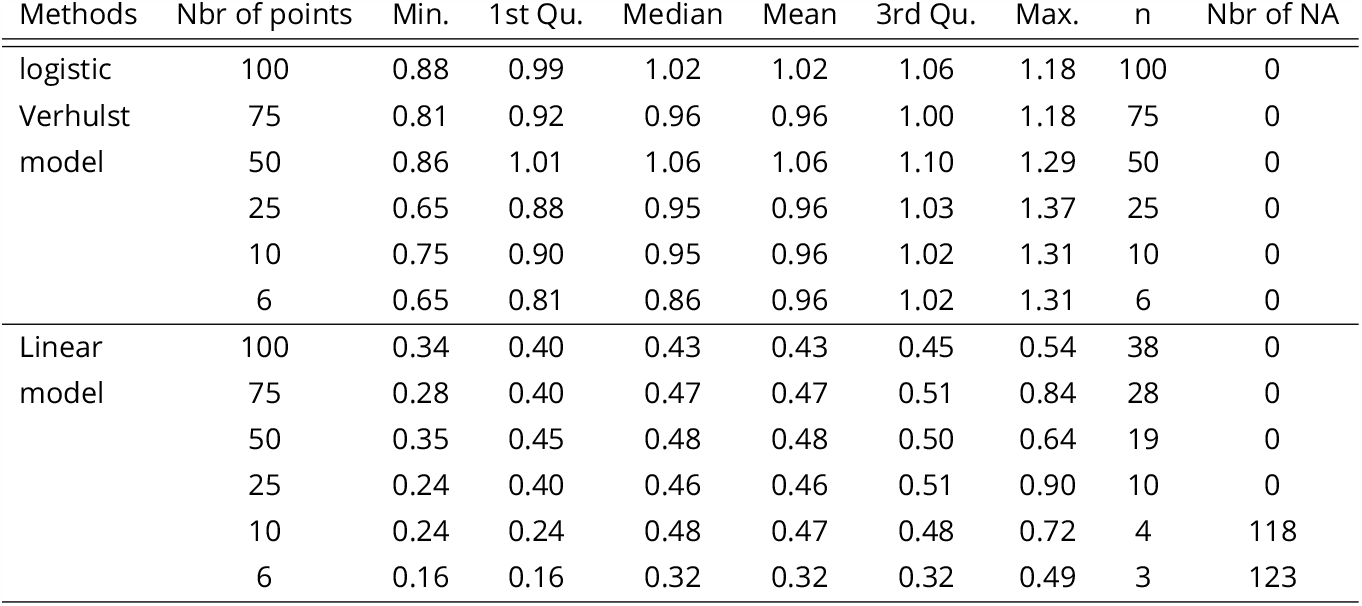
Statistical summary of growth rates estimated by logistic Verhulst model and linear model obtained after 1000 bootstrap simulations. Min. is the minimum growth rate, 1stQu. is the 1st quartile, 3rdQu. is the 3rd quartile and Max. is the maximum, *n* is the number of points used by each method to estimate the growth rate and Nbr of NA is the number of simulations that did not result into a growth rate estimate.

| Methods | Nbr of points | Min. | 1st Qu. | Median | Mean | 3rd Qu. | Max. | n | Nbr of NA |
| --- | --- | --- | --- | --- | --- | --- | --- | --- | --- |
| logistic Verhulst model | 100 | 0.88 | 0.99 | 1.02 | 1.02 | 1.06 | 1.18 | 100 | 0 |
|  | 75 | 0.81 | 0.92 | 0.96 | 0.96 | 1.00 | 1.18 | 75 | 0 |
|  | 50 | 0.86 | 1.01 | 1.06 | 1.06 | 1.10 | 1.29 | 50 | 0 |
|  | 25 | 0.65 | 0.88 | 0.95 | 0.96 | 1.03 | 1.37 | 25 | 0 |
|  | 10 | 0.75 | 0.90 | 0.95 | 0.96 | 1.02 | 1.31 | 10 | 0 |
|  | 6 | 0.65 | 0.81 | 0.86 | 0.96 | 1.02 | 1.31 | 6 | 0 |
| Linear model | 100 | 0.34 | 0.40 | 0.43 | 0.43 | 0.45 | 0.54 | 38 | 0 |
|  | 75 | 0.28 | 0.40 | 0.47 | 0.47 | 0.51 | 0.84 | 28 | 0 |
|  | 50 | 0.35 | 0.45 | 0.48 | 0.48 | 0.50 | 0.64 | 19 | 0 |
|  | 25 | 0.24 | 0.40 | 0.46 | 0.46 | 0.51 | 0.90 | 10 | 0 |
|  | 10 | 0.24 | 0.24 | 0.48 | 0.47 | 0.48 | 0.72 | 4 | 118 |
|  | 6 | 0.16 | 0.16 | 0.32 | 0.32 | 0.32 | 0.49 | 3 | 123 |

**Figure 6.**
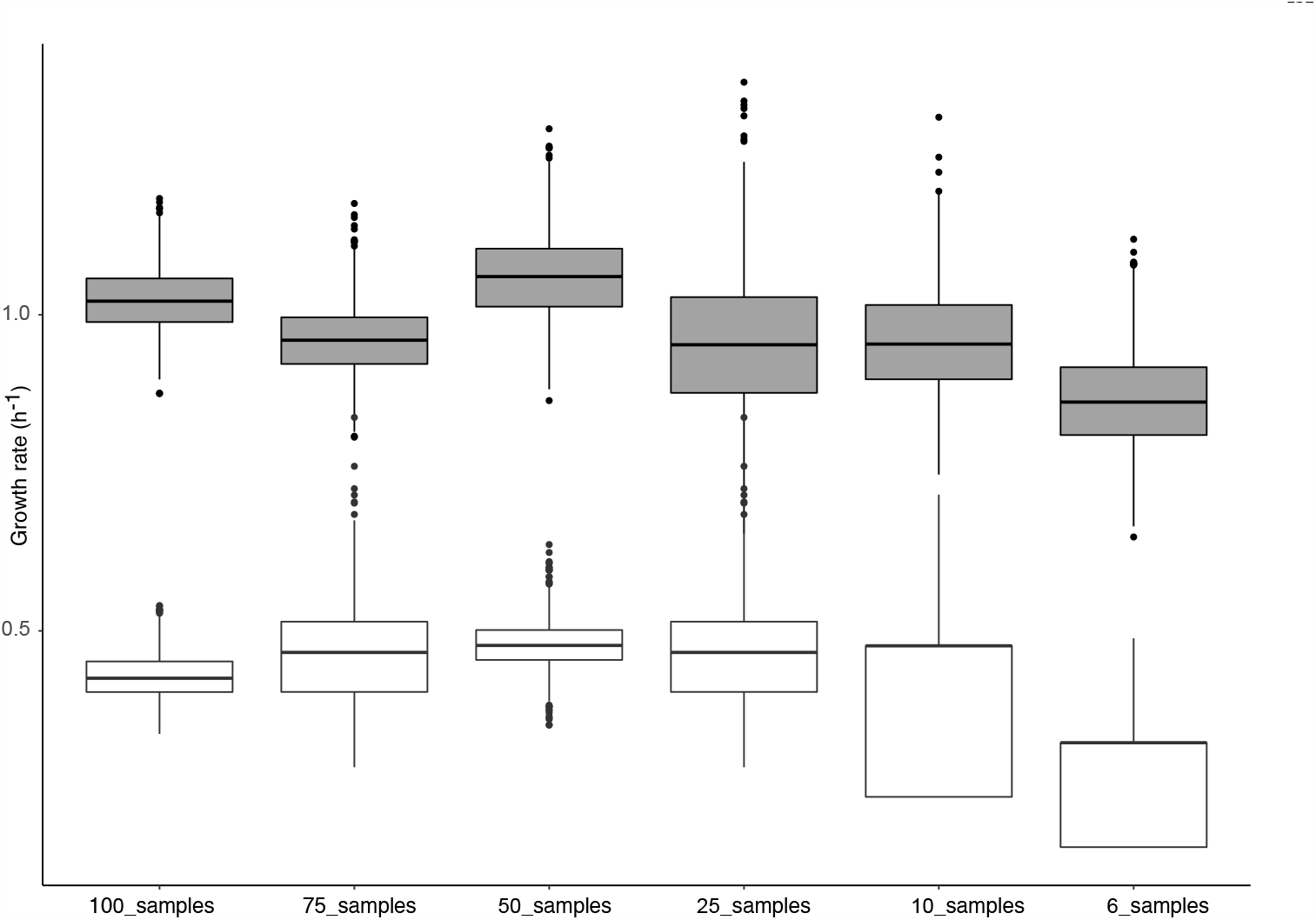
Distribution of the *μ*_*max*_ estimated for 1000 simulations for each curve. The black bar represents the median value. The gray box plots represent the distribution of *μ*_*max*_ estimated by the logistic Verhulst method and the white box plots represent *μ*_*max*_ estimated by the linear model in the exponential phase after linearization of the data.

In Table 1, it is interesting to see that with a limited number of points, of either 10 or 6, the linear model can not be applied in more than a hundred of simulations per curve (number of NA respectively 118 and 123). Such result appears when the number of points used into the model (n in Table 1) is very small in the exponential phase. In microbiology such case is frequent, especially for experiments in which numerous variables are sampled at the same time, and can not be measured quickly enough to catch the exponential phase, or for overnight bacterial growth.

Looking at the literature, 17 growth curves have been collected (Table 2) from various environments including freshwater, deep-sea, hydrothermal vents or sediments. This table compares the estimation of growth rate *μ*_*max*_ using: logistic Verhulst model, linear model model and the *μ*_*max*_ value provided by author. The main goal of this computation is to apply logistic Verhulst model on existing datasets from the literature. All the data are extracted from the growth curves represented in each article, then the two models are applied. Firstly, we can note that growth rate provided by the authors are very close to those estimated by linear model, as expected, since this method is widely used in microbiology. However, using logistic Verhulst model, the *μ*_*max*_ is higher than those estimated by linear model and provided by authors for 88% (15 times over the 17 reported growth curves) of curves. Moreover, the logistic Verhulst model allows estimating two other parameters, the beginning of the exponential growth and the maximum biomass. The beginning of the exponential growth is a critical parameter in food industry (Zwietering et al., 1990). In deep-sea environments Martini et al. (2013), the use of the growth rate and the maximum biomass (K parameter in this model) allow computing cross-coefficient that turned out to be a paramount tool to determine optimal growth conditions for a deep sea strain of luminous bacteria. Ultimately, the modeling approach allows transforming discrete data into a continuous function. This last point, is a critical step to associate the bacterial growth with other high frequency variables measured simultaneously (i.e.: oxygen consumption (Garel, Bonin, et al., 2019), light emission (Al Ali et al., 2010)).

**Table 2.**
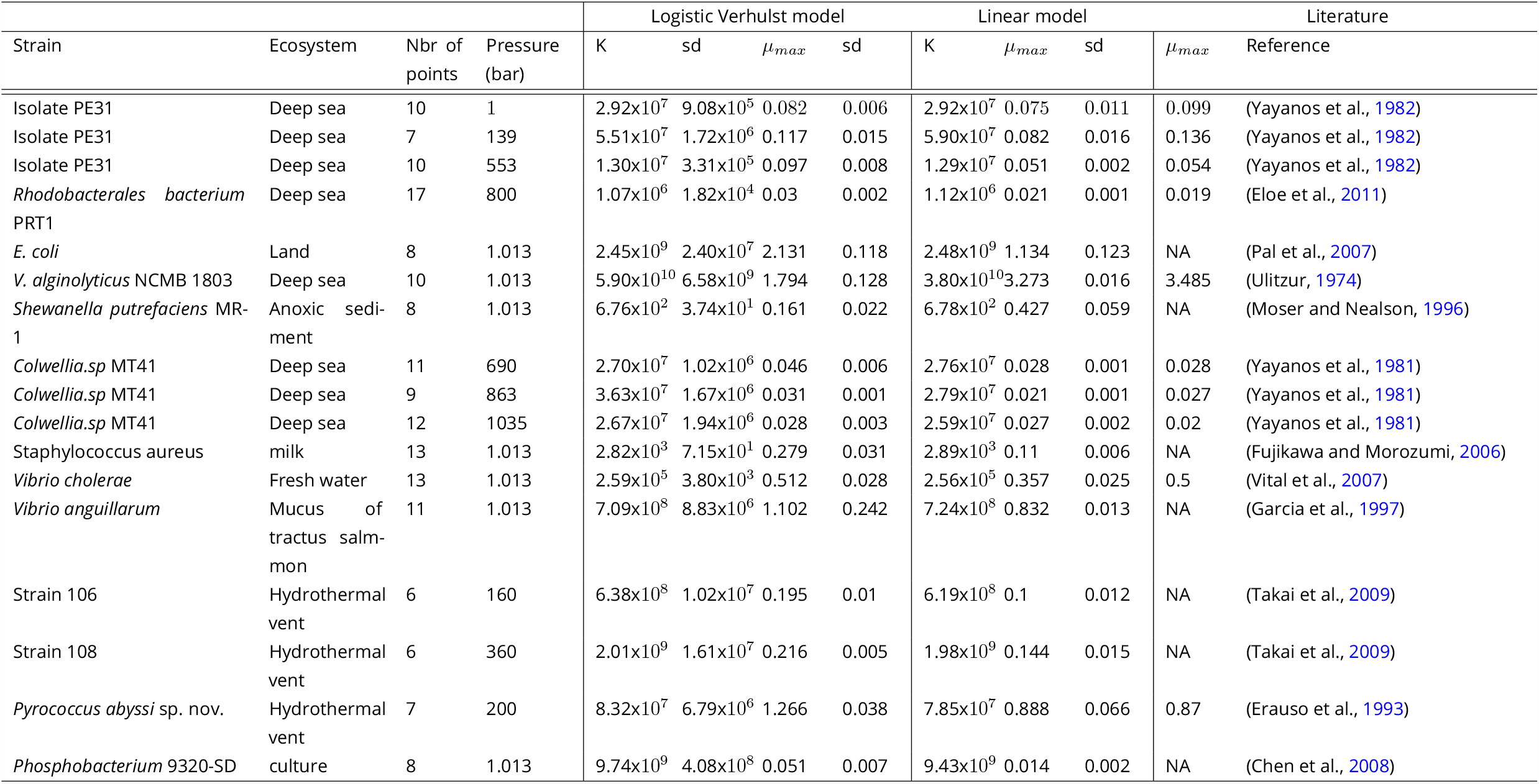
Synthesis of 17 bacterial strains from various aquatic ecosystems. Growth parameters are estimated by Logistic Verhulst model and Linear model. K is the maximum cells density, sd is the standard deviation and *μ*_*max*_ is the growth rate

| Strain | Ecosystem | Nbr of points | Pressure (bar) | Logistic Verhulst model |  |  |  | Linear model |  |  | Literature |  |
| --- | --- | --- | --- | --- | --- | --- | --- | --- | --- | --- | --- | --- |
| | | | | K | sd | $\mu_{max}$ | sd | K | $\mu_{max}$ | sd | $\mu_{max}$ | Reference |
| Isolate PE31 | Deep sea | 10 | 1 | 2.92x10<sup>7</sup> | 9.08x10<sup>5</sup> | 0.082 | 0.006 | 2.92x10<sup>7</sup> | 0.075 | 0.011 | 0.099 | (Yayanos et al., 1982) |
| Isolate PE31 | Deep sea | 7 | 139 | 5.51x10<sup>7</sup> | 1.72x10<sup>6</sup> | 0.117 | 0.015 | 5.90x10<sup>7</sup> | 0.082 | 0.016 | 0.136 | (Yayanos et al., 1982) |
| Isolate PE31 | Deep sea | 10 | 553 | 1.30x10<sup>7</sup> | 3.31x10<sup>5</sup> | 0.097 | 0.008 | 1.29x10<sup>7</sup> | 0.051 | 0.002 | 0.054 | (Yayanos et al., 1982) |
| *Rhodobacterales bacterium* PRT1 | Deep sea | 17 | 800 | 1.07x10<sup>6</sup> | 1.82x10<sup>4</sup> | 0.03 | 0.002 | 1.12x10<sup>6</sup> | 0.021 | 0.001 | 0.019 | (Eloe et al., 2011) |
| *E. coli* | Land | 8 | 1.013 | 2.45x10<sup>9</sup> | 2.40x10<sup>7</sup> | 2.131 | 0.118 | 2.48x10<sup>9</sup> | 1.134 | 0.123 | NA | (Pal et al., 2007) |
| *V. alginolyticus* NCMB 1803 | Deep sea | 10 | 1.013 | 5.90x10<sup>10</sup> | 6.58x10<sup>9</sup> | 1.794 | 0.128 | 3.80x10<sup>10</sup> | 3.273 | 0.016 | 3.485 | (Ulitzur, 1974) |
| *Shewanella putrefaciens* MR-1 | Anoxic sedi-ment | 8 | 1.013 | 6.76x10<sup>2</sup> | 3.74x10<sup>1</sup> | 0.161 | 0.022 | 6.78x10<sup>2</sup> | 0.427 | 0.059 | NA | (Moser and Nealson, 1996) |
| *Colwellia.sp* MT41 | Deep sea | 11 | 690 | 2.70x10<sup>7</sup> | 1.02x10<sup>6</sup> | 0.046 | 0.006 | 2.76x10<sup>7</sup> | 0.028 | 0.001 | 0.028 | (Yayanos et al., 1981) |
| *Colwellia.sp* MT41 | Deep sea | 9 | 863 | 3.63x10<sup>7</sup> | 1.67x10<sup>6</sup> | 0.031 | 0.001 | 2.79x10<sup>7</sup> | 0.021 | 0.001 | 0.027 | (Yayanos et al., 1981) |
| *Colwellia.sp* MT41 | Deep sea | 12 | 1035 | 2.67x10<sup>7</sup> | 1.94x10<sup>6</sup> | 0.028 | 0.003 | 2.59x10<sup>7</sup> | 0.027 | 0.002 | 0.02 | (Yayanos et al., 1981) |
| *Staphylococcus aureus* | milk | 13 | 1.013 | 2.82x10<sup>3</sup> | 7.15x10<sup>1</sup> | 0.279 | 0.031 | 2.89x10<sup>3</sup> | 0.11 | 0.006 | NA | (Fujikawa and Morozumi, 2006) |
| *Vibrio cholerae* | Fresh water | 13 | 1.013 | 2.59x10<sup>5</sup> | 3.80x10<sup>3</sup> | 0.512 | 0.028 | 2.56x10<sup>5</sup> | 0.357 | 0.025 | 0.5 | (Vital et al., 2007) |
| *Vibrio anguillarum* | Mucus of tractus salm-mon | 11 | 1.013 | 7.09x10<sup>8</sup> | 8.83x10<sup>6</sup> | 1.102 | 0.242 | 7.24x10<sup>8</sup> | 0.832 | 0.013 | NA | (Garcia et al., 1997) |
| Strain 106 | Hydrothermal vent | 6 | 160 | 6.38x10<sup>8</sup> | 1.02x10<sup>7</sup> | 0.195 | 0.01 | 6.19x10<sup>8</sup> | 0.1 | 0.012 | NA | (Takai et al., 2009) |
| Strain 108 | Hydrothermal vent | 6 | 360 | 2.01x10<sup>9</sup> | 1.61x10<sup>7</sup> | 0.216 | 0.005 | 1.98x10<sup>9</sup> | 0.144 | 0.015 | NA | (Takai et al., 2009) |
| *Pyrococcus abyssi* sp. nov. | Hydrothermal vent | 7 | 200 | 8.32x10<sup>7</sup> | 6.79x10<sup>6</sup> | 1.266 | 0.038 | 7.85x10<sup>7</sup> | 0.888 | 0.066 | 0.87 | (Erauso et al., 1993) |
| *Phosphobacterium* 9320-SD | culture | 8 | 1.013 | 9.74x10<sup>9</sup> | 4.08x10<sup>8</sup> | 0.051 | 0.007 | 9.43x10<sup>9</sup> | 0.014 | 0.002 | NA | (Chen et al., 2008) |

From a statistical point of view, comparing these two modelling approaches on the same dataset we observe differences. On Figure 7, both approaches are applied on the growth curve of the bacterial strain *Shewanella putrefaciens*. The logistic Verhulst model (Figure 7A) estimates a growth rate of 0.161±0.02 *h*^−1^ while, according to data selected to estimate the slope of the line in the exponential phase, the grow rate can vary from 0.38 à 0.55 *h*^−1^ (Figure 7B).

**Figure 7.**
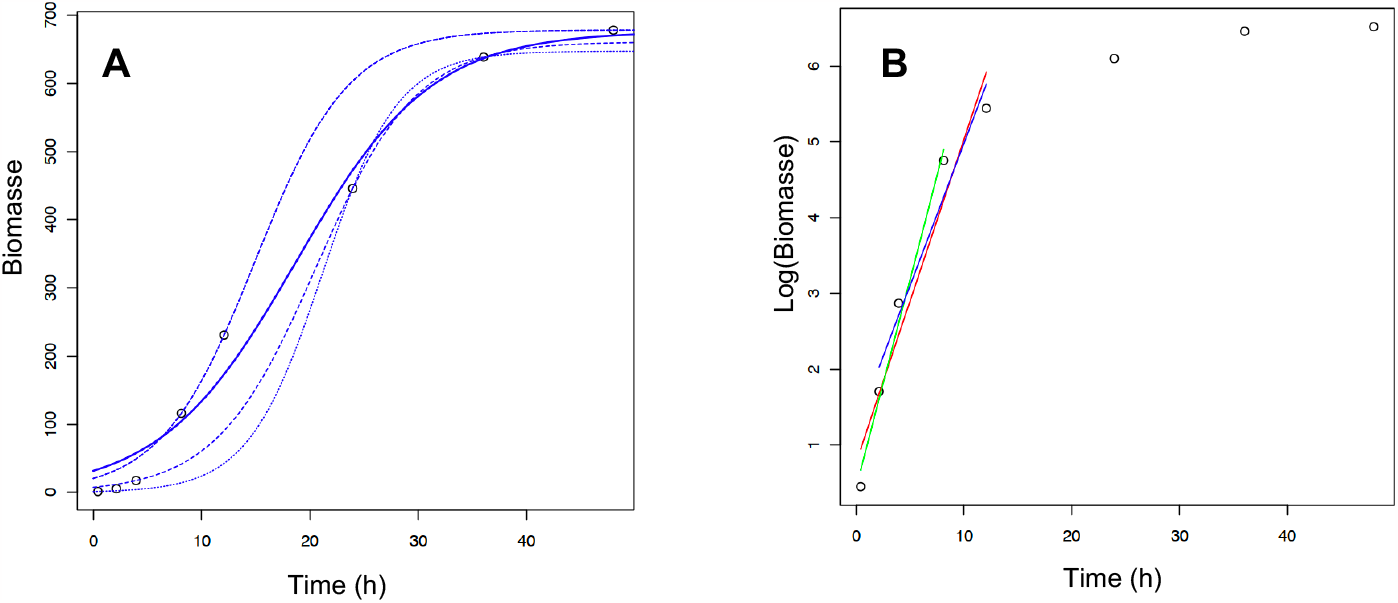
Comparison of growth rate estimations using the logistic Verhulst model and the linear model. A, the growth rate is estimated with the logistic Verhulst model. B, the growth rate is estimated based on the linear model applied on three different combination of points describing the exponential phase (red, blue and green lines).

The distribution of growth rate is less variable with the logistic Verhulst model than the linear model (Figure 6), even if the confidence interval can be large with the logistic Verhulst model. Moreover, the main advantage of the modeling approach is the objectivity and lack of user bias. Indeed, for the logistic Verhulst model all points are taken into account while for the linear model *μ*_*max*_ the linear part of the curve is estimated subjectively to determine the slope (i.e. the growth rate) of this curve section (Zwietering et al., 1990).

## Conclusions

Pluridisciplinarity is essential in order to solve scientific problems in various fields of life science such as ecology or microbiology. However, biologists needs to have access to statistical or modelling tools already developed by mathematicians. In this work we give access to a web application dedicated to the modelling and computing of growth parameters, essential for microbiologists, without deep mathematical knowledge. Statistics are also available in order to estimate the model efficiency.

## Acknowledgements

Authors would like to thank Maurice Libes (OSU Pytheas) for his help to deploy the code on a public repository. Authors would like to thank Marie Roumagnac (MIO), Chloé Baumas (MIO) and Emelyne Vidal (ICMCB) for the beta testing of the web application.

## References

Al Ali B, M Garel, P Cuny, JC Miquel, T Toubal, A Robert, and C Tamburini (2010). Luminous bacteria in the deepsea waters near the ANTARES underwater neutrino telescope (Mediterranean Sea). Chemistry and Ecology 26, 57–72. ISSN: 0275-7540. 10.1080/02757540903513766.

Chen Z, S Ma, and L Liu (2008). Studies on phosphorus solubilizing activity of a strain of phosphobacteria isolated from chestnut type soil in China. Bioresource Technology 99, 6702–6707. ISSN: 09608524. 10.1016/j.biortech.2007.03.064.

Eichinger M, S Kooijman, R Sempere, D Lefevre, G Gregori, B Charriere, and JC Poggiale (2009). Consumption and release of dissolved organic carbon by marine bacteria in a pulsed-substrate environment: from experiments to modelling. Aquatic Microbial Ecology 56, 41–54. ISSN: 0948-3055. 10.3354/ame01312.

Eichinger M, JC Poggiale, and R Sempere (2011). Toward a mechanistic approach to modeling bacterial DOC pathways: a review. Microbial carbon pump in the ocean. Science booklet, Supplement to Science, 66–68.

Eloe EA, F Malfatti, J Gutierrez, K Hardy, WE Schmidt, K Pogliano, J Pogliano, F Azam, and DH Bartlett (2011). Isolation and characterization of a psychropiezophilic alphaproteobacterium. Applied and environmental microbiology 77, 8145–53. ISSN: 1098-5336. 10.1128/AEM.05204-11.

Erauso G, AL Reysenbach, A Godfroy, JR Meunier, B Crump, F Partensky, J Baross, V Marteinsson, G Barbier, N Pace, and D Prieur (1993). Pyrococcus abyssi sp. nov., a new hyperthermophilic archaeon isolated from a deep-sea hydrothermal vent. Archives of Microbiology 160. ISSN: 0302-8933. 10.1007/BF00252219.

Fujikawa H and S Morozumi (2006). Modeling Staphylococcus aureus growth and enterotoxin production in milk. FOOD MICROBIOLOGY 23, 260–267. 10.1016/j.fm.2005.04.005.

Garcia T, K Otto, S Kjelleberg, and DR Nelson (1997). Growth of Vibrio anguillarum in salmon intestinal mucus. Applied and Environmental Microbiology 63, 1034–1039. ISSN: 00992240. 10.1128/aem.63.3.1034-1039.1997.

Garel M, P Bonin, S Martini, S Guasco, M Roumagnac, N Bhairy, F Armougom, and C Tamburini (2019). Pressure-Retaining Sampler and High-Pressure Systems to Study Deep-Sea Microbes Under in situ Conditions. Frontiers in Microbiology 10, 453. ISSN: 1664-302X 10.3389/fmicb.2019.00453.

Garel M, C Panagiotopoulos, M Boutrif, D Repeta, R Sempere, C Santinelli, B Charriere, D Nerini, JC Poggiale, and C Tamburini (2021). Contrasting degradation rates of natural dissolved organic carbon by deep-sea prokaryotes under stratified water masses and deep-water convection conditions in the NW Mediterranean Sea. Marine Chemistry 231, 103932. ISSN: 03044203. 10.1016/j.marchem.2021.103932.

Gompertz B (1825). XXIV. On the nature of the function expressive of the law of human mortality, and on a new mode of determining the value of life contingencies. In a letter to Francis Baily, Esq. F. R. S. &c. Philosophical Transactions of the Royal Society of London 115, 513–583. ISSN: 0261-0523. 10.1098/rstl.1825.0026.

Kempes CP, S Dutkiewicz, and MJ Follows (2012). Growth, metabolic partitioning, and the size of microorganisms. Proceedings of the National Academy of Sciences 109, 495–500. ISSN: 0027-8424. 10.1073/pnas.1115585109. eprint: https://www.pnas.org/content/109/2/495.full.pdf.

Lobry JR, JP Flandrois, G Carret, and A Pave (1992). Monod’s bacterial growth model revisited. Bulletin of Mathematical Biology 54, 117–122. ISSN: 15229602. 10.1007/BF02458623.

Martini S, B Al Ali, M Garel, D Nerini, V Grossi, M Pacton, L Casalot, P Cuny, and C Tamburini (2013). Effects of Hydrostatic Pressure on Growth and Luminescence of a Moderately-Piezophilic Luminous Bacteria Photobacterium phosphoreum ANT-2200. PLoS ONE 8. Ed. by Driks A, e66580. ISSN: 1932-6203. 10.1371/journal.pone.0066580.

Monod J (1949). The Growth of Bacterial Cultures. Annual Review of Microbiology 3, 371–394. ISSN: 0066-4227. 10.1146/annurev.mi.03.100149.002103.

Moser DP and KH Nealson (1996). Growth of the Facultative Anaerobe Shewanella putrefaciens by Elemental Sulfur Reduction Downloaded from. Tech. rep. 6, pp. 2100–2105.

Pal S, YK Tak, and JM Song (2007). Does the antibacterial activity of silver nanoparticles depend on the shape of the nanoparticle? A study of the gram-negative bacterium Escherichia coli. Applied and Environmental Microbiology 73, 1712–1720. ISSN: 00992240. 10.1128/AEM.02218-06.

R Core Team (2017). R: A language and environment for statistical computing.

Richards FJ (1959). A flexible growth function for empirical use. Journal of Experimental Botany 10, 290–301. ISSN: 00220957. 10.1093/jxb/10.2.290.

Schnute J (1981). A versatile growth model with statistically stable parameters. Canadian Journal of Fisheries and Aquatic Sciences 38, 1128–1140. ISSN: 12057533. 10.1139/f81-153.

Scott M and T Hwa (2011). Bacterial growth laws and their applications. 10.1016/j.copbio.2011.04.014.

Stannard CJ, AP Williams, and PA Gibbs (1985). Temperature/growth relationships for psychrotrophic food-spoilage bacteria. Food Microbiology 2, 115–122. ISSN: 10959998. 10.1016/S0740-0020(85)80004-6.

Takai K M Miyazaki, H Hirayama, S Nakagawa, J Querellou, and A Godfroy (2009). Isolation and physiological characterization of two novel, piezophilic, thermophilic chemolithoautotrophs from a deep-sea hydrothermal vent chimney. Environmental Microbiology 11, 1983–1997. ISSN: 14622912. 10.1111/j.1462-2920.2009.01921.x.

Ulitzur S (1974). Vibrio parahaemolyticus and Vibrio alginolyticus: Short generation-time marine bacteria. Microbial Ecology 1, 127–135. ISSN: 00953628. 10.1007/BF02512384.

Verhulst P (1845). Recherches mathématiques sur la loi d’accroissement de la population.; Recherches mathématiques sur la loi d’accroissement de la population. Mém. Acad. r. Sci. Lett. Belg. 18, 1–38.

Verhulst P(1847). Deuxième mémoire sur la loi d’accroissement de la population. Mémoires de l’Académie Royale des Sciences, des Lettres et des Beaux-Arts de Belgique 20, 1–32.

Verhulst P(1938). Notice sur la loi que la population suit dans son accroissement. Quetelet Quetelet 1, 113–121.

Vital M, HP Füchslin, F Hammes, and T Egli (2007). Growth of Vibrio cholerae O1 Ogawa Eltor in freshwater. Microbiology 153, 1993–2001. ISSN: 13500872. 10.1099/mic.0.2006/005173-0.

Yayanos AA, AS Dietz, and R Van Boxtel (1981). Obligately barophilic bacterium from the Mariana trench. Proceedings of the National Academy of Sciences 78, 5212–5215. ISSN: 0027-8424. 10.1073/pnas.78.8.5212.

Yayanos AA, AS Dietz, and R Van Boxtel(1982). Dependence of Reproduction Rate on Pressure as a Hallmark of Deep-Sea Bacteria. Applied and Environmental Microbiology 44, 1356–1361. ISSN: 0099-2240. 10.1128/aem.44.6.1356-1361.1982.

Zwietering MH, I Jongenburger, FM Rombouts, and K van ‘t Riet (1990). Modeling of the bacterial growth curve. Applied and environmental microbiology 56, 1875–81.

